# H3-K27M mutation alters the dynamics of human hematopoietic stem cells and delays erythroid differentiation

**DOI:** 10.64898/2026.09.14.751581

**Authors:** Mia Brunetti, Hassan Dakik, Fatemeh Beigmohammadi, Kolja Eppert, Morgan Craig

**Affiliations:** Département de Mathématiques et de Statistiques, Université de Montréal, Montréal, Canada; Sainte-Justine University Hospital Research Center, Montréal, Canada; Department of Pediatrics, McGill University, Montréal, Canada; Research Institute of the McGill University Health Centre, Canada

## Abstract

Acute myeloid leukemia (AML) is an aggressive blood cancer driven by genetic and epigenetic alterations that disrupt normal hematopoiesis. Among these, the histone H3-K27M mutation, originally identified in pediatric high-grade gliomas, reshapes gene repression programs by reducing global H3-K27 trimethylation. Although rare, H3-K27M mutations have been detected in preleukemic hematopoietic stem cells (HSCs) of AML patients, suggesting its role in early leukemogenesis and identifying it as a promising therapeutic target.

Here, we investigated how H3-K27M alters hematopoiesis using a longitudinal xenotransplantation mouse model of human hematopoietic stem and progenitor cells carrying either wild-type H3 or H3-K27M. To quantify hematopoietic dynamics, we developed a collection of mathematical models representing alternative lineage hierarchies and fit them to our longitudinal experimental data. Using information criteria, we identified the model that best depicts blood dynamics in each condition. Our results show that H3-K27M promotes HSC proliferation and differentiation and erythroid commitment while reducing multipotent progenitor self-renewal potential and lymphoid commitment, ultimately increasing blood cell numbers despite impairing erythroid maturation. Mathematical modelling inferred the progenitor-level alterations, which were not readily evident from the blood cell count alone. H3-K27M also induced distinct kinetics in human HSC1 and HSC2 subpopulations that were not observed in wild-type H3 controls. Together, our results provide the first quantitative framework to reveal how H3-K27M influences the hematopoietic hierarchy roadmap and alters blood cell dynamics in preleukemia.

**Key Points:**

- Mathematical modelling identifies H3-K27M–induced changes in hematopoietic kinetics that underlie impaired erythroid maturation.
- H3-K27M amplifies kinetic differences between HSC1 and HSC2, with HSC1 driving mutant HSC expansion.

## Introduction

Acute myeloid leukemia (AML) is an aggressive and heterogeneous hematological malignancy that develops through the sequential acquisition of genetic alterations in hematopoietic stem and progenitor cells (HSPCs) [1-3]. These altered progenitors enter a preleukemic state before progressing to leukemic clones that expand into blasts with an impaired differentiation. AML outcomes remain poor despite current standard-of-care therapy [4], which primarily debulk these highly proliferating cells from the bone marrow and blood [5].

In AML and other myeloid malignancies, the disruption of epigenetic mechanisms driven by mutations in histone genes and in genes encoding regulators such as *EZH2* and *ASXL1* contribute to leukemogenesis at the preleukemic phase [6-9]. Indeed, the H3-K27M mutation, first discovered in pediatric high-grade gliomas [10, 11], was recently identified in patients with AML [8]. This lysine 27 to methionine substitution on the histone H3 tail domain inhibits the enzymatic activity of the polycomb repressive complex 2 (PRC2), decreasing the overall repressive H3-K27 trimethylation marks vital for transcriptional control affecting cell fate and differentiation [12, 13]. *In vitro* and *in vivo* murine and human xenotransplantation studies have shown that H3-K27M increases the frequency of functional hematopoietic stem cells (HSCs), alters myeloid differentiation, and enhances leukemic aggressiveness [8, 14]. Additionally, high variant allele frequencies of H3-K27M in both primary AML and remission samples suggest that this mutation arises in major preleukemic HSC clones [8], making it a promising therapeutic target. Nevertheless, it remains unclear how H3-K27M modifies the dynamics of blood cells during preleukemia.

Mathematical modelling has been used to characterize healthy and abnormal blood cell production, helping to establish mechanisms of altered blood cell dynamics [15-24]. Pioneering work by Busch et al. defined the kinetics of hematopoiesis at homeostasis by modelling the progression of an inducible label throughout the blood hierarchy in both developing and adult mice [25]. Similar frameworks have identified alternative differentiation pathways to the classical hematopoietic hierarchy in healthy blood [26] and determine whether clonal hematopoiesis alters the blood’s hierarchical organization [27]. However, these later studies were parameterized to human *in vitro* data and were applied to slow-progressing disorders, unlike AML. Thus, the *in vivo* dynamics of aggressive preleukemia, particularly in human cells, remain largely unexplored.

To better understand the role of H3-K27M in early leukemogenesis, we performed a longitudinal *in vivo* xenotransplantation study using human CD34+CD38-HSPCs transduced with H3-K27M or wild-type H3 to investigate how this mutation influences HSC proliferation and hematopoietic lineage fate over time. To further elucidate how H3-K27M reshapes hematopoietic dynamics in human HSPCs, we developed multiple mathematical models representing alternative lineage hierarchies. After parameterizing these models to our longitudinal data from human HSPCs, we selected the most parsimonious models to determine the most likely hematopoietic hierarchy given observed H3-K27M and wild-type H3 blood dynamics. Our results show that, in this human-cell system, H3-K27M drives the increase of all blood cell compartments through HSC expansion relative to wild-type, resulting in a decreased lymphoid lineage commitment of multipotent progenitors (MPPs) and a latent blockage in erythroid differentiation. Further, our mathematical model quantified the dynamics underlying these alterations and revealed the inferred structural changes in hematopoietic stem cell organization induced by H3-K27M, providing a quantitative framework to understand the progression of preleukemia to AML.

## Methods

### Time-course xenograft experiment

Cord blood samples were collected from male and female donors with informed consent. To reduce the influence of variability between donors and to avoid results being driven by effects from outliers, all samples were pooled before isolating human CD34+ hematopoietic stem and progenitor cells (HSPCs). CD34^+^CD38^−^ cells were sorted and transduced with either histone H3-WT control or H3-K27M mutant lentiviral vectors, as described previously [8]. Vectors contained a hybrid bidirectional promoter (SFFV/minimal CMV) driving the expression of both GFP and the insert gene. Xenotransplantation took place four days after transduction and mice received irradiation with 2.1 Gy 24 hours before HSPC transplantation. The cells were injected into the right femurs of female NOD.Cg-PrkdscidII2rgtm1Wjl/SzJ (NSG) mice, aged to 11 and 23 weeks, which equate to around 20,000 CD34^+^CD38^−^ cells at the time of transduction. Intrafemoral injections were performed under isoflurane anesthesia, with buprenorphine (0.1 mg/kg) administered for analgesia. Control mice received human cord blood-driven HSPCs overexpressing the histone H3 wild-type gene. Both control (H3-WT) and experimental mice transplanted with human cells overexpressing histone H3-K27M (H3-K27M) were kept in parallel under identical experimental conditions and maintained on a regular diet. Mice were first purchased from The Jackson Laboratory, then bred in-house. Ten H3-WT and H3-K27M mice were euthanized at 4, 8, 12 and 16 weeks via cervical dislocation following isoflurane anesthesia. The number of samples per time point was determined based on prior experience with the model [8]. The animal-use protocol was approved by McGill University and the Research Institute of McGill University Health Centre.

### Flow cytometry and cell count quantification

Bone marrow cells were harvested from pooled femurs, tibias and pelvises by mechanical crushing of the bones, washed, and resuspended in 500 µl of PBS containing 2% of COSMIC calf serum (CCS; HyClone, SH30087.03). Absolute cell counts and population frequencies were quantified by flow cytometry on a BD LSRFortessa (BD Biosciences) equipped with a high-throughput sampler (HTS), using three independent antibody panels targeting established CD markers: 1) a global hematopoietic panel to assess engraftment and lineage output; 2) a differentiation panel to assess mature myeloid, erythroid and lymphoid populations; and 3) a progenitor panel to assess immunophenotypic HSCs and downstream myeloid and lymphoid progenitors. The gating strategy for each panel is shown in **Supplementary Figure S1**. Prior to staining with the progenitor panel, human HSPCs were enriched and mouse cells depleted using the EasySepTM Human Progenitor Cell Enrichment Kit with platelet depletion (STEMCELL Technologies, #19356) and the EasySepTM Mouse/Human Chimera Isolation Kit (STEMCELL Technologies, #19849).

The global panel was used to quantify the absolute number of total transduced (GFP^+^) cells and of CD45+, CD34^+^, as well as CD33^+^ myeloid and CD19^+^ lymphoid cells within the GFP^+^ compartment. The differentiation panel was used to quantify transduced CD71^+^GlyA^+^ and CD71^−^GlyA^+^ erythroid populations. The progenitor panel was used to quantify common myeloid progenitors (CMPs; CD45^+^ CD34^+^ CD38^+^ CD7^−^ CD10^−^ CD135^+^ CD45RA^−^), granulocyte-monocyte progenitors (GMPs; CD45^+^ CD34^+^ CD38^+^ CD7^−^ CD10^−^ CD135^+^ CD45RA^+^), megakaryocyte–erythroid progenitors (MEPs; CD45^+^ CD34^+^ CD38^+^ CD7^−^ CD10^−^ CD135^−^ CD45RA^−^), multi-lymphoid progenitors (MLPs; CD45^+^ CD34^+^ CD38^−^ CD45RA^+^), multipotent progenitors (MPPs; CD45^+^ CD34^+^ CD38^−^ CD45RA^−^ CD90^−^ CD49f^−^), HSC1 (CD45^+^ CD34^+^ CD38^−^ CD45RA^−^ CD90^+^ CD49f^+^), and HSC2 (CD45^+^ CD34^+^ CD38^−^ CD45RA^−^ CD90^−^ CD49f^+^). Analysis was done in flowJo software (v10). Percentages of progenitor populations were first determined within the CD34^+^ compartment, and absolute cell numbers were calculated using CD34^+^ cell counts obtained from the global panel.

Panels were composed of antibodies, listed in the **Supplementary Methods**, with corresponding fluorophore, clone and dilution information. All antibodies were purchased from BioLegend unless otherwise specified. Sytox blue (1:1000 dilution) was used for viability staining (Life Technologies; S34857).

### Mathematical models of hematopoietic hierarchy

We used a bottom-up approach [28] to build our mathematical models of hematopoiesis that consisted of multiple systems of ordinary differential equations, with each model variable representing a different compartment in the differentiation pathway of blood cells (**Figure 1; Supplementary Methods, Eqs. (1)-(18)**).

**Figure 1.**
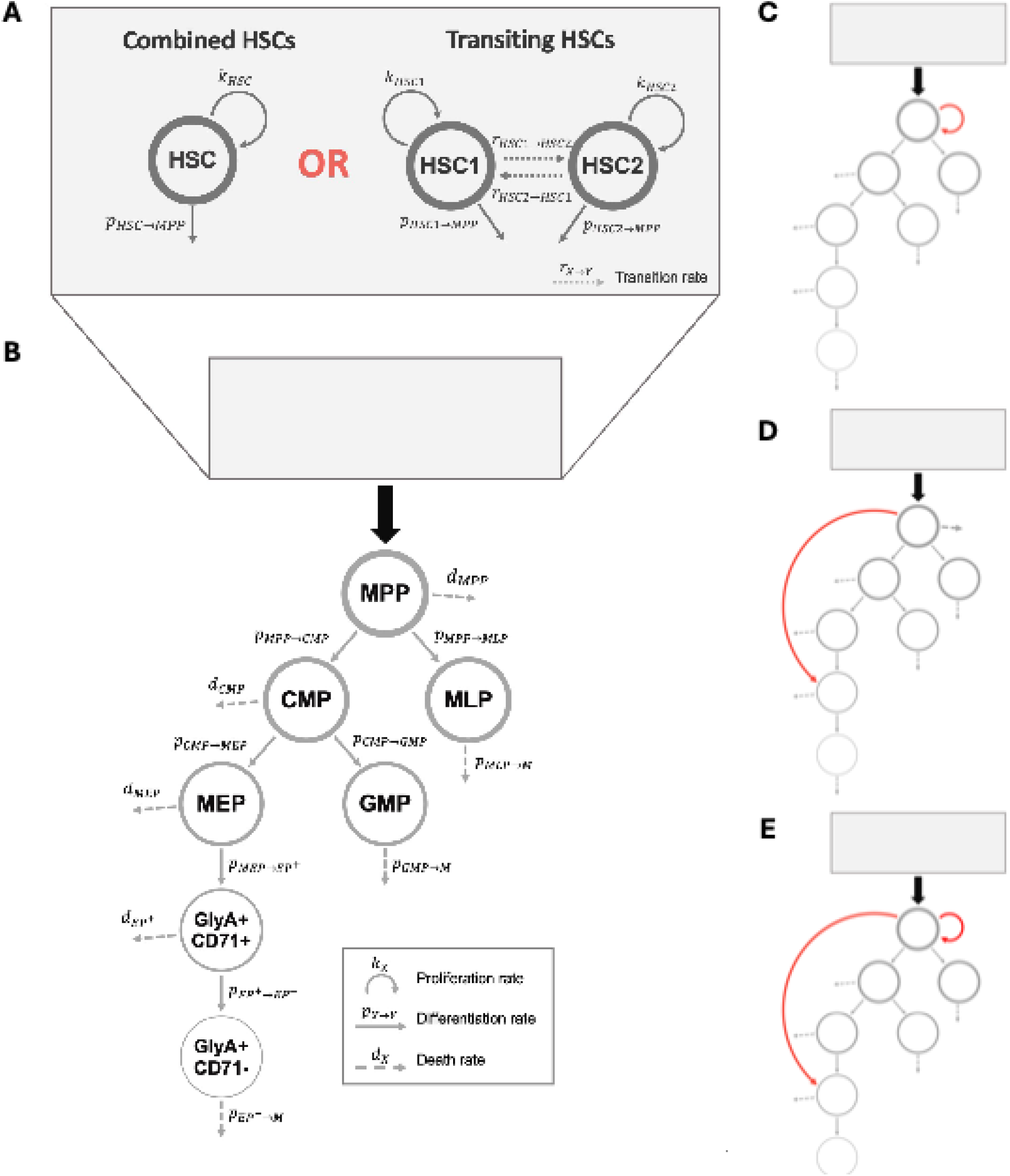
Plausible mathematical models of the hematopoietic hierarchy. Schematics of the eight hematopoietic hierarchies and differentiation pathways tested. Each model describes dynamics in the hematopoietic stem cells (HSC), five progenitors (MPP, MLP, CMP, GMP, and MEP), and two GlyA+ erythroid precursors with varying levels of maturity (EP+ and EP™) compartments. More mature, unmeasured progenitor compartments are identified as M. A) Models describing the organization of the HSC compartment. In the Combined HSCs model, HSC1 and HSC2 are considered as a single stem cell population with the same dynamics. In the Transiting HSCs model, HSC1 and HSC2 are considered as distinct populations in which stem cells can transit from one population to the other. All eight models correspond to the two HSC structures in A combined with each of the models describing the rest of hematopoiesis in B, C, D, and E. B) Classical model of the hematopoietic hierarchy [34-36]. C) Same model as in B but with MPP self-renewal. D) Same model as in B but with a direct differentiation pathway from MPP to EP+ (MEP bypass). E) Same model as in D but with MPP self-renewal. Circular arrows: proliferation. Solid arrows: differentiation into populations included in the model. Dashed arrows: death/net outflow (i.e., death and differentiation into unmodelled populations M). Dotted arrows: transition between populations.

At the top of the hierarchy, HSCs can self-renew and differentiate into multipotent progenitors (MPPs) (**Figure 1A; Supplementary Methods, Eqs. (1), (3), and (4)**). Two populations of hematopoietic stem cells, HSC1 and HSC2, exist in humans, mainly differing in their expression of CD90 surface cell markers and their fraction of repopulating cells [29]. While it is known that CD90+ cells produce CD90cells, *in vivo* experiments have shown that CD90-cell transplants can generate CD90+ within the bone marrow of NSG mice [29, 30]. To better understand the relationship between HSC1 and HSC2, we constructed two models denoted as the Combined HSCs model and the Transiting HSCs model (**Figure 1A**). In the Combined HSCs model, we assume that the hematopoietic stem cell compartment includes both the HSC1 and HSC2 populations and has a unique self-renewal rate and differentiation rate to MPPs (**Supplementary Methods, Eq. (1)**), whereas HSC1 and HSC2 compartments were modelled separately, each with their own self-renewal rate and differentiation rate into MPPs, in the Transiting HSCs model. In the Transiting HSCs model, cells can move between the HSC1 and HSC2 compartments at different rates (**Supplementary Methods, Eqs. (3) and (4)**).

For the MPP compartment, we considered two cases: 1) without self-renewal where MPPs are simply removed by natural death (**Figure 1B; Supplementary Methods, Eqs. (2), (5), (12), and (13)**) and 2) where they can self-renew (**Figure 1C; Supplementary Methods, Eqs. (15)-(18)**). In all hematopoietic organizational structures, we considered MPPs to generate the two main blood lineages by differentiating, with distinct rates, into multi-lymphoid progenitors (MLPs) and common myeloid progenitors (CMPs) (**Figure 1B-E; Supplementary Methods, Eqs. (2), (5), (12), (13), (15)-(18)**). Downstream, CMPs differentiate into both granulocyte-macrophage progenitors (GMPs) and megakaryocyte-erythroid progenitors (MEPs) at different rates (**Supplementary Methods, Eq. (6)**). The model further describes the sequential differentiation of MEPs into early CD71+ GlyA+ erythroid precursors (EP+) and more mature CD71™GlyA+ erythroid precursors (EP™). No other compartment follows MLPs, GMPs, and EP+, because no data from downstream populations were used. For MLPs, GMPs and EP+, we define the net outflow rate as the sum of the differentiation rate into more matured progenitors (M), which are not modelled, and the death rate (**Figure 1B-E; Supplementary Methods, Eqs. (6)-(11)**). Otherwise, except for compartments with self-renewal capacities (i.e., the HSCs and, in some models, the MPPs), all other cell compartments have specific death rates.

Lastly, multiple studies have shown alternative pathways for erythroid differentiation in which cells higher in the hierarchy can differentiate directly into erythroid precursors [31-33]. Thus, to consider a direct differentiation pathway from MPPs to the early EP+ cells (**Figure 1D-E**), we modified the classical hematopoietic hierarchy [34-36] by adding a rate of differentiation from the MPP to the EP+ compartment (**Supplementary Methods, Eqs. (12)-(14)**).

### Parameter estimation

After assessing structural identifiability (see **Supplementary Methods; Table S1**), we fit each candidate ordinary differential equation model to both the H3-WT and H3-K27M xenotransplantation data. We used adaptive simulated annealing (ASA) [37, 38], a probabilistic global optimization technique, to minimize the weighted residual sum of squares (WRSS) between the model and the data (**Supplementary Methods, Eq. (19)**). To find the set of best fit parameters, we performed 200 ASA iterations, each initialized to a randomly chosen set of initial guesses sampled from a uniform distribution given by the theoretical limits for each parameter. The best fit parameters were those for which the WRSS was the lowest among the 200 ASA runs. For detailed explanations of the parameter estimation process, see **Supplementary Methods**.

### Distributions and credible intervals of best fit estimates

For each best fit parameter of the selected model for H3-WT and H3-K27M, we estimated their distribution using trajectory-matching random-walk feasibility sampling (TM-RWFS) [39]. This algorithm is a model- and data-based approach that captures the variability of complex biological systems. Parameter distributions are generated by constraining the model’s trajectories between the upper and lower bounds of available time-series data. We forced the trajectories to lie between the minimum and maximum values of all data points, except at t = 56 days of the H3-K27M HSC2 cell counts. We relaxed the conditions for this time point, since the best fit value for HSC2 in the H3-K27M dynamics exceeds the value of the highest observation. Parameters were sampled from normal distributions, which we initialized with a mean equal to the best fit parameter value and a chosen standard deviation of 0.05. For each condition, we restricted the exploration of the parameter space with theoretical limits like those used for parameter estimation (see **Parameter Estimation**) to remain within the local region of the best fit values in the optimization space. We ran the algorithm until 1000 trajectories were accepted (**Figure S2**). The distributions of the parameters of the best fit models are shown in **Figure 3A** (for shared parameters) and **Figure S3**.

We then calculated the 95% credible intervals of the best fit parameters for both H3-WT and H3-K27M dynamics by calculating the 95% highest density interval (HDI) of each parameter’s distributions [40] (see **Table S2**). Similarly, we used the HDI from the trajectory distributions at every simulated time point for all blood cell populations for model solutions for both H3-WT and H3-K27M (**Figure 2**). For further information, see **Supplementary Methods**.

**Figure 2.**
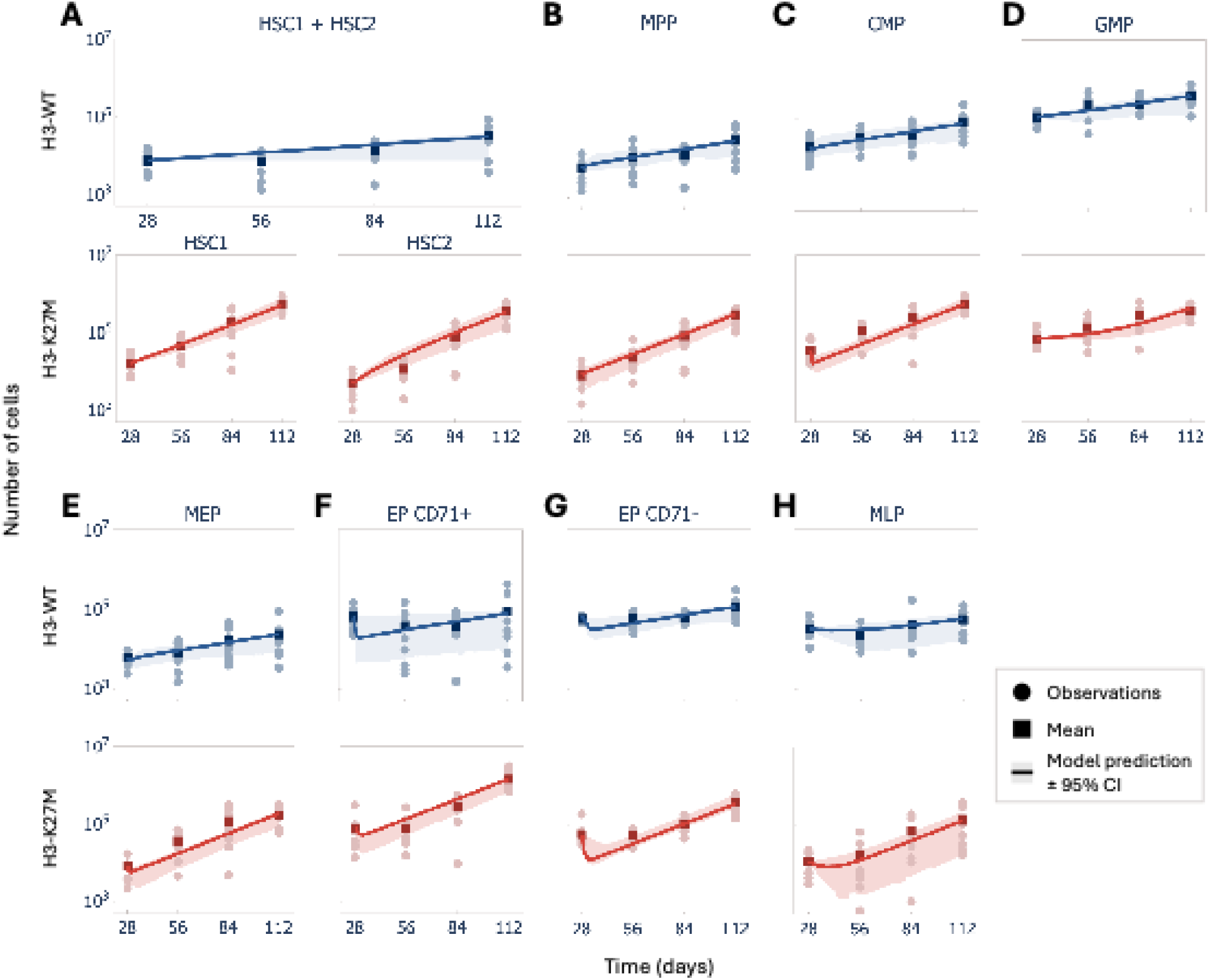
H3-K27M accelerates the preleukemic expansion of HSCs. Model C predicted growth curves fit to the number of H3-WT (blue; Combined HSCs structure) or H3-K27M (red; Transiting HSCs structure) cells across hematopoietic populations. **A)** HSC1+HSC2 combined for H3-WT and HSC1 and HSC2 for H3-K27M. **B)** MPP. **C)** CMP. **D)** GMP. **E)** MEP. **F)** EP CD71+. **G)** EP CD71™. **H)** MLP. Solid circles indicate individual mouse observations; squares indicate means; solid lines indicate model prediction; shaded areas indicate predicted 95% credible intervals.

### Relative change in model parameters

To compare the values of the fitted parameters between the H3-WT and H3-K27M model, we calculated the relative change of shared parameters for H3-K27M relative to H3-WT. Let p_WT_ and p_K27M_ be the H3-WT and H3-K27M best fit values for the shared parameter p, respectively. The relative change is described with the following equation:

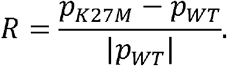

### Statistical analysis

Distributions of common parameters between the selected models were compared using the Mann-Whitney U test [41], with rank-biserial correlation (r_rb_) used as an effect size measure. Detailed procedure and application are provided in the **Supplementary Methods**.

## Results

### Classical differentiation structure best describes healthy and H3-K27M hematopoiesis

To determine the structure of hematopoietic differentiation in healthy and H3-K27M mutated human blood cells and to assess whether we can uncover a separate erythroid differentiation pathway from the blood cell counts, we performed a systematic comparison of our various mathematical models (**Figure 1**). For this, we used the corrected version of the Akaike information criterion (AICc), a statistical tool for model selection that ranks models depending on the goodness of fit and complexity [42, 43]. We fit all models to our experimental results of H3-WT and H3-K27M longitudinal dynamics (**Figure 2; Tables S2-S6**) and ranked them according to ΔAICc (**Table 1**), i.e., the difference between the AICc value for each model and that of the most parsimonious model (**Supplementary Methods, Eqs. (23)-(24)**).

**Table 1.** Hematopoietic hierarchy model selection results for H3-WT and H3-K27M experimental data. Most plausible models are highlighted in grey. ΔAICc: Difference between the AICc value of each model and most parsimonious model. WRSS: weighted residual sum of squares.

| Models | Combined HSCs |  |  |  | Transiting HSCs |  |  |  |
| --- | --- | --- | --- | --- | --- | --- | --- | --- |
|  | H3-WT |  | H3-K27M |  | H3-WT |  | H3-K27M |  |
| | $\Delta AICc$ | WRSS | $\Delta AICc$ | WRSS | $\Delta AICc$ | WRSS | $\Delta AICc$ | WRSS |
| **B** | 0.0 | 221.4 | 13.1 | 176.2 | 3.9 | 252.7 | 2.2 | 197.1 |
| **C** | 0.0 | 221.5 | 8.2 | 173.2 | 3.0 | 252.0 | 0.0 | 195.7 |
| **D** | 25.6 | 240.1 | 16.8 | 177.1 | 5.8 | 252.4 | 4.4 | 197.0 |
| **E** | 10.7 | 228.1 | 12.0 | 174.2 | 6.1 | 252.7 | 4.3 | 196.9 |

Regardless of the mathematical structure of the HSC compartment(s), models B and C received the strongest AICc support for H3-WT (**Table 1**; **Figure 1B-C**). However, model B required HSC to MPP differentiation rates higher than 5.458 days^-1^ (**Tables S3 and S4**), which is substantially larger than reported experimental estimates [44-46]. Therefore, model B was excluded on biological-plausibility grounds. Model C was also the best-supported mathematical model for H3-K27M (**Table 1**). Hence, all subsequent comparisons between H3-WT and H3-K27M dynamics were performed using model C, with parameters estimated separately for each genotype. These results indicate that H3-K27M alters the inferred kinetics within the selected hematopoietic structures. They do not, however, by themselves establish that the mutation changes the underlying hierarchy.

Furthermore, this approach did not favour models D and E (**Figure 1D-E**) that incorporate the alternative erythroid differentiation pathway (MEP bypass). Thus, given our data, a direct differentiation pathway between MPPs and EP+ does not improve the ability of the model to explain the hematopoietic dynamics of both H3-WT and H3-K27M cells.

### H3-K27M enhances the distinction between HSC1 and HSC2 kinetics

Our AICc ranking approach best supports the Combined HSCs and the Transiting HSCs formulations for H3-WT and H3-K27M, respectively (**Table 1; Figure 1A**). These rankings do not demonstrate a genotype-specific modification in HSC organization, as they were determined independently for each genotype. Rather, this result suggests that HSC1 and HSC2 subsets, while distinct, are less distinguishable under healthy hematopoiesis, potentially because of their relatively low proliferation and cellular division rates. The increased proliferative capacity of HSCs carrying the H3-K27M mutation may amplify kinetic differences between the HSC subpopulations, allowing for HSC1 and HSC2 to be differentiated from one another in the model.

### H3-K27M promotes the intrinsic growth of HSCs and increases overall blood cell counts

The data reveals that by day 112, all cell populations, except for GMPs and MLP, harbouring the H3-K27M mutation, undergo a more significant expansion than those with the wild-type H3 (**Figure 2; Figure S3; Supplementary Methods**). This trend is also evident in the selected models (**Figure S4A**). The largest difference occurs in the HSCs: using the models’ prediction, we calculated that, together, the HSC1 and HSC2 subpopulations carrying H3-K27M expand to 28.06 times the size of the wild-type HSC (Combined HSCs) at day 112 (**Figure S4B**). This trend was more variable throughout the time course of our experiments (i.e., from days 0 to 112) (**Figure S4B**). Indeed, normalized (see **Supplementary Methods**) H3-K27M populations of HSCs, MPPs, CMPs, MEPs, and EP+ were predicted to be consistently higher than corresponding H3-WT hematopoietic populations, whereas GMPs, EP™, and MLPs H3-K27M mutant cell counts remained below those for equivalent H3-WT populations (**Figure S4B**).

We sought to better understand these growth differences by directly comparing the estimated kinetic rates for each cell compartment (i.e., HSC/HSC1 and HSC2, MPP, CMP, GMP, MEP, EP+, EP™, and MLP). Indeed, using model C with the Combined HSCs structure for H3-WT and the Transiting HSCs structure for H3-K27M, we found increased HSC expansion in H3-K27M transplanted mice compared with H3-WT (net proliferation rate of the total HSC population (HSC1 + HSC2) was estimated to be a_HSC_ = k_HSC_ - p_HSC➔MPP_ = 0.017 days^-1^ in H3-WT_HSC1 HSC2_ versus a = 0.031 days^-1^ and a = 0.011 days^-1^ for HSC1 and HSC2, respectively, in the H3-K27M model; **Table S2**). Further, because the H3-K27M HSC population is described as the sum of two exponential terms, its long-term dynamics are dominated by the larger growth rate, a_HSC1_. Since a_HSC1_ exceeds a_HSC_, the model predicts a higher long-term expansion in H3-K27M HSCs versus the wild-type population and that is mainly driven by the HSC1 subpopulation. Overall, our results show that while all blood cell populations expand from their initial value in both the wild-type and the mutant cases, the growth of hematopoietic populations is much higher in H3-K27M cells (**Figure S4A**).

### H3-K27M disrupts erythroid differentiation and other downstream hematopoietic dynamics

Lastly, we sought to understand how altered kinetic rates in H3-K27M mutations drive the dynamic changes described above. For this, we began by looking for changes in the estimated parameter distributions for both the wild-type and mutant (**Figure 3A; Figure S5**). Mann-Whitney U tests of each pair of distributions showed statistically significant differences (all p < 1 × 10^−16^), indicating that the dynamics induced by H3-K27M are different from those of the wild-type (**Figure 3A**; **Table S7**). Specifically, we found the differentiation rates p_CMP→MEP_ and p_MEP→EP+_, the net outflow rates p_MLP→M_ and p_EP-→M_, and the death rate d_CMP_ were to have a positive rank-biserial correlation (r_rb_ > 0), with r_rb_ values ranging from 0.361 (moderately separated, for) to 0.769 (highly separated, for), indicating higher values for H3-K27M versus wild-type. Conversely, the self-renewal rate k_MPP_, differentiation rates p_MPP→MLP_, p_MPP→CMP_, p_CMP→ MP_, p_EP+→EP-_, the net outflow rate p_MP→M_, and the death rates d_MEP_ and d_EP+_ were all reduced in H3-K27M relative to H3-WT. In the latter cases, distributions ranged from completely separated (r_rb_ = -1, for k_MPP_) to moderately separated (r_rb_ = -0.461, for d). The direction of these H3-K27M-induced parameter changes is summarized schematically in **Figure 3C**.

**Figure 3.**
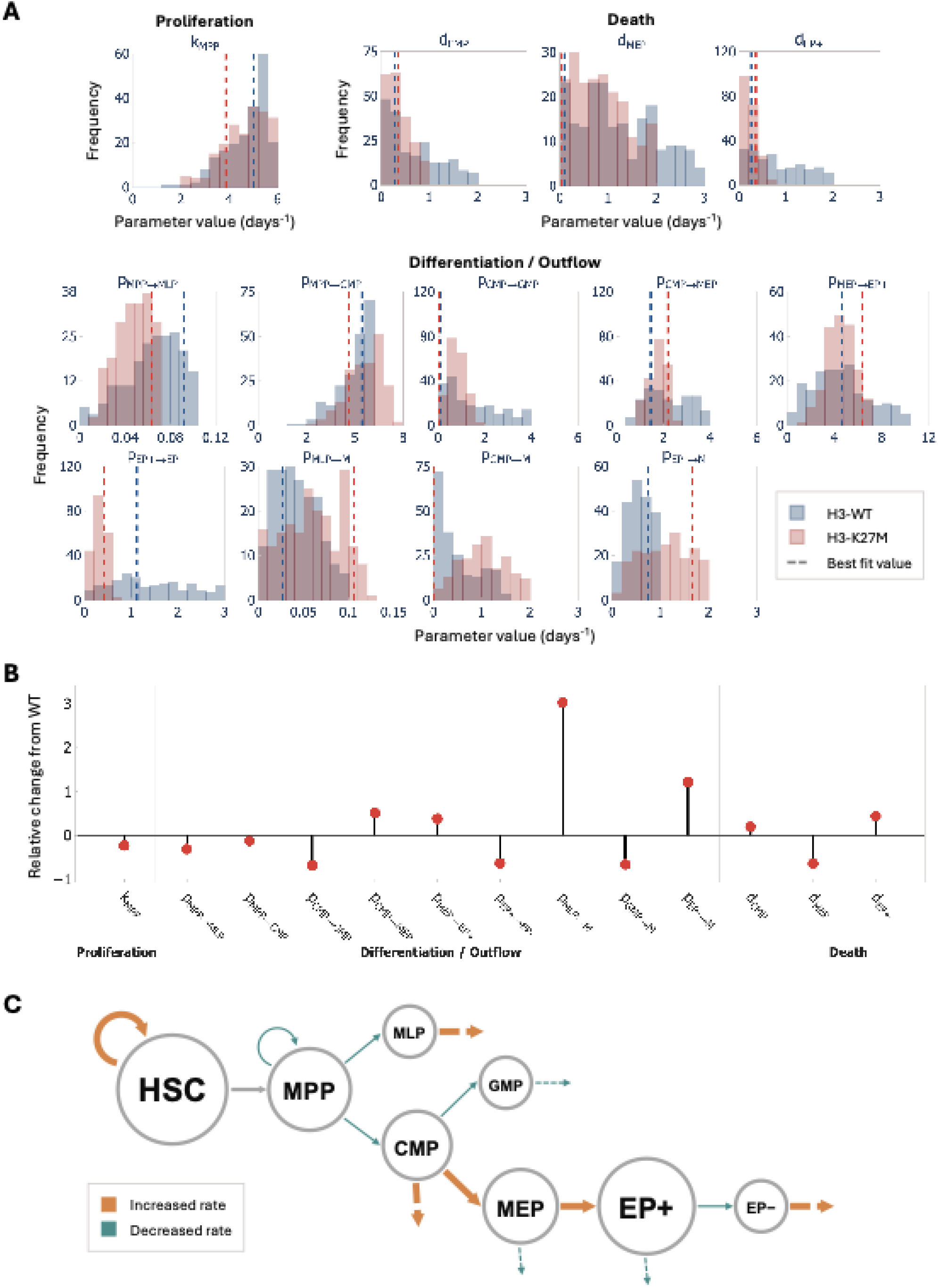
H3-K27M drives myeloid bias while reducing erythroid differentiation. Comparison of model kinetics that are shared between best fit model C, with the Combined and Transiting HSC structure for H3-WT and H3-K27M, respectively. **A)** Accepted distributions of parameter estimates based on 1,000 accepted trajectories using TM-RWSF [39] (see Supplementary Information). Each subplot displays the overlapping distributions of a shared parameter from the model C fit to H3-WT (blue) and H3-K27M (red) data. Vertical dashed line: best fit value. **B)** Relative change of best fit values of shared parameters for H3-K27M relative to H3-WT. **C)** Schematic summarizing how H3-K27M influences hematopoietic compartment size and kinetics. Relative increase in cell count in a hematopoietic population compared to H3-WT is represented by a larger circle: GMP, MLP, and EP™ cell counts are approximately equivalent in both H3-WT and H3-K27M. Thick orange arrows: increased hematopoietic rates in H3-K27M. Thin turquoise arrows: decreased hematopoietic rates in H3-K27M.

Based on these differences, we next compared estimated best fit parameter values to check for magnitudes of differences by calculating the relative change in shared model parameters between H3-WT and H3-K27M (**Figure 3B**). The largest change was found in the net outflow rate from the MLP compartment, p_MLP→M_, which increased by 3.0-fold for the H3-K27M case, followed by the net outflow rate of EP™ cells, p_EP-→M_, which was increased 1.21-fold. These findings reveal compensatory mechanisms in altered H3-K27M hematopoiesis. Indeed, the increased expansion of upstream compartments, the higher rate of HSC proliferation, and the reduced rate of MPP differentiation increase MLP levels, albeit to a lesser extent than for upstream HSC and MPP populations. Consequently, the net outflow rate of MLPs, p_MLP→M_, must increase to balance this expansion. Similarly, despite the marked expansion of MEP and EP+ populations, the compartment of EP™ expands more modestly, translating to an increase in p_EP-→M_. **Figure 3C** integrates these rate changes across the selected hematopoietic model C.

## Discussion

Acute myeloid leukemia is a hard-to-treat and aggressive disease that evolves through the acquisition of multiple mutations during the preleukemic phase [3]. Previous studies have detected patterns of mutations in preleukemia wherein epigenetic mutations appear to occur early in mutational cascades [7, 9]. How such early mutations alter hematopoietic dynamics during preleukemia, resulting in speeding up or slowing these evolutionary cascades, is an open question. The histone mutation H3-K27M has been previously identified in patients with AML but it has not been extensively studied in the context of preleukemia. Our *in vivo* human xenograft model allows us to study the progression of normal and H3-K27M mutated human hematopoiesis within living tissues. To characterize changes to the structure of the hematopoietic hierarchy and blood cell production dynamics induced by H3-K27M, we combined our mouse xenotransplantation data with a biologically informed mathematical model selection framework, ultimately identifying the underlying kinetics driving H3-K27M dysregulated hematopoiesis.

Overall, our model selection framework favoured the classical model of hematopoiesis without an MEP bypass for both H3-WT and H3-K27M, consistent with previous studies [26, 27]. However, differences emerged within the HSC compartment, revealing the interplay between CD34+CD38-CD45RA-CD90+CD49f+ (HSC1) and CD34+CD38-CD45RA-CD90-CD49f+ (HSC2) subsets. In particular, the selection of the Combined HSCs model C suggests that HSC cellular division occurs too infrequently in the H3-WT case to distinguish these two sub-populations. By promoting the expansion rate of HSCs, the H3-K27M mutation accentuates and quantifies the differences in dynamics between HSC1 and HSC2, with the HSC1 sub-population driving mutant HSC expansion by dividing more frequently than HSC2, as predicted by its higher rates of self-renewal and differentiation into MPPs. This agrees with experimental observations that describe HSC1 as containing a higher fraction of repopulating cells than HSC2 [29].

Our results also identify how H3-K27M deregulates hematopoietic dynamics. In addition to increased HSC proliferation and differentiation, our modelling predicted an overall decrease in MPP differentiation and lymphoid commitment. Interestingly, the self-renewal rate of MPPs was also predicted to be decreased in H3-K27M hematopoiesis relative to H3-WT, reflecting a reduced contribution of MPP self-renewal and a greater contribution of the HSC compartment to hematopoietic maintenance. Downstream of MPPs, our model predicted an increased CMP differentiation toward the MEP lineage, followed by an increase in erythroid production from MEPs coupled with an impaired maturation of the erythroid precursors in the case of H3-K27M. These two changes—the reduced MPP self-renewal rate and increased differentiation of CMPs into MEPs—are particularly noteworthy, as they are not readily apparent from our blood cell count data alone. While all blood cell populations with H3-K27M generally expand and reach counts above those of the wild-type cells, these impaired dynamics provide a mechanistic explanation for the massive expansion of CD71+GlyA+ erythroid precursors.

While we found our models to closely describe our data, we did not evaluate all plausible differentiation pathways that may exist within the hematopoietic hierarchy. Our model selection results should also be interpreted with the caveat that identifying alternative differentiation pathways may require measurements of additional hematopoietic cell populations or complementary data beyond blood cell counts. Indeed, the lack of support for models with, e.g., an alternative erythroid differentiation pathway (MEP bypass) does not exclude the possibility that such a pathway exists biologically. Rather, it implies that it was less supported by our data. Moreover, some model parameters would not have been structurally identifiable, given the data used for fitting, and addressing this identifiability issue would require additional experimental measurements. Lastly, our models do not account for extrinsic factors influencing hematopoietic dynamics, nor do they integrate interactions between healthy and H3-K27M mutated blood cells. Extending our models to include these mechanisms represents an important future direction.

Defining the hierarchical structure and the proliferation and differentiation kinetics of blood cell populations is essential to improve our understanding of hematopoiesis and the dysregulation that arises prior to and during hematological malignancies. In turn, this allows us to better predict the response of blood diseases to treatment and to define specific therapeutic targets. Our computational work resulted in parameterized mechanistic mathematical models of both normal and H3-K27M mutated hematopoiesis that identify the hematopoietic populations and rates most affected by this preleukemic mutation. Together, these findings provide a quantitative framework for investigating the mechanisms leading to the development of AML and guiding future experimental and therapeutic studies.

## Acknowledgments

This work was supported, in part, by the Canadian Institutes of Health Research [183641]; the Canada Research Chairs Program in Computational Immunology; the Fonds de recherche du Québec – Santé [https://doi.org/10.69777/329621]; the Cole Foundation; and the New Directions in Research Competition from the Montreal Children’s Hospital Foundation and the Research institute of the McGill University Health Centre.

## Authorship Contributions

MB: Conceptualization, methodology, software, formal analysis, visualization, writing – original draft.

HD: Conceptualization, methodology, investigation, formal analysis, visualization, writing – original draft and review & editing.

FB: Methodology, software, writing – review & editing.

KE: Conceptualization, methodology, resources, formal analysis, writing – review & editing, supervision.

MC: Conceptualization, methodology, formal analysis, writing – review & editing, supervision.

