## Supplemental Data for "H3-K27M mutation alters the dynamics of human hematopoietic stem cells and delays erythroid differentiation"

### Supplementary Methods

**Antibodies in flow cytometry panels**

*Global panel:* CD45 (AlexaFluor 700; clone 2D1; 1:100), CD34 (APC-Cy7; clone 581; 1:100), CD38 (PE; clone HB-7; 1:1000), CD33 (APC; clone WM53; 1:300), CD19 (BV711; clone HIB19; 1:100) and CD235a (PerCP-Cy5.5; clone HI264; 1:100).

*Differentiation panel:* CD45 (AlexaFluor 700; clone 2D1; 1:100), CD33 (APC; clone WM53; 1:300), CD14 (PE:Dazzle 594; clone HCD14; 1:250), CD11b (BV650; clone ICRF44; 1:100), CD235a (PerCP-Cy5.5; clone HI264; 1:100), CD71 (PE; clone MA712; 1:1000; BD Biosciences), CD41 (PE-Cy7; clone HIP8; 1:100), CD3 (BV605; clone SK7; 1:250), CD19 (BV711; clone HIB19; 1:100) and CD56 (PE-Cy5; clone hCD56; 1:250).

*Progenitors panel:* CD45 (AlexaFluor 700; clone 2D1; 1:100), CD34 (APC-Cy7; clone 581; 1:100), CD38 (PE; clone HB-7; 1:1000), CD7 (PE-Cy7; clone CD7-6B7; 1:500), CD10 (Pe-Cy5; clone HI10a; 1:500), CD45RA (BV650; clone HI100; 1:100), FLT3 (PerCP-Cy5.5; clone BV10A4H2; 1:100), CD90 (BV605; clone 5E10; 1:100) and CD49f (APC; clone GoH3; 1:500).

**Mathematical models of hematopoiesis**

We constructed eight different models (B, C, D, and E; see **Figure 1** and **Methods** in the Main Text) consisting of systems of ordinary differential equations to describe the dynamics of HSC (HSC1/HSC2), MPP, CMP, GMP, MEP, EP+, and EP- cells. Each model follows the differentiation, proliferation, and death of these cell populations during hematopoiesis. The *Combined HSCs* models describe the dynamics of cell populations in the set $S_{C}\in$ [HSC, MPP, CMP, GMP, MEP, EP+, EP-] whereas the *Transiting HSCs* models differentiate between HSC1 and HSC2 subsets to describe cell populations in $S_{T}\in$ [HSC1, HSC2 MPP, CMP, GMP, MEP, EP+, EP-].

Let $k_{i}$ be the self-renewal rate of cell type $i$, $p_{i\to j}$ the differentiation rate from cell type $i$ to $j$, $d_{i}$ the natural death rate of cell type $i$, and $r_{i\to j}$ the transition rate from cell type $i$ to $j$. For cell populations that can self-renew, such as HSCs and, in some models, MPPs, we used a net proliferation rate that equals the difference between their self-renewal rate and their natural death rate. Similarly, the outflux from MLPs, GMPs, and CD70- erythroid progenitors is described using a net differentiation rate corresponding to the sum of their differentiation rate and their natural death rate (see Structural Identifiability section below).

Then, the dynamics of HSC and MPP populations in the *Combined HSCs* models are given by:

$$\frac{dHSC}{dt}=\left( k_{HSC}-p_{HSC\to MPP} \right)HSC$$

(1)

$$\frac{dMPP}{dt}=p_{HSC\to MPP}HSC-\left( d_{MPP}+p_{MPP\to CMP}+p_{MPP\to MLP} \right)MPP$$

(2)

while those for the *Transiting HSCs* model are given by:

$$\frac{dHSC\text{1}}{dt}=\left( k_{HSC1}-p_{HSC1\to MPP}-r_{HSC1\to HSC2} \right)HSC\text{1}+r_{HSC2\to HSC1}HSC\text{2}$$

(3)

$$\frac{dHSC\text{2}}{dt}=\left( k_{HSC2}-p_{HSC2\to MPP}-r_{HSC2\to HSC1} \right)HSC\text{2}+r_{HSC1\to HSC2}HSC\text{1}$$

(4)

$$\frac{dMPP}{dt}=p_{HSC1\to MPP}HSC\text{1}+p_{HSC2\to MPP}HSC\text{2}-\left( d_{MPP}+p_{MPP\to CMP}+p_{MPP\to MLP} \right)MPP.$$

(5)

Downstream of the MPPs, we have:

$$\frac{dCMP}{dt}=p_{MPP\to CMP}MPP-\left( d_{CMP}+p_{CMP\to MEP}+p_{CMP\to GMP} \right)CMP$$

(6)

$$\frac{dGMP}{dt}=p_{CMP\to GMP}CMP-p_{GMP\to M}GMP$$

(7)

$$\frac{dMEP}{dt}=p_{CMP\to MEP}CMP-\left( d_{MEP}+p_{MEP\to{EP}^{+}} \right)MEP$$

(8)

$$\frac{d{EP}^{+}}{dt}=p_{MEP\to{EP}^{+}}MEP-\left( d_{{EP}^{+}}+p_{{EP}^{+}\to{EP}^{-}} \right){EP}^{+}$$

(9)

$$\frac{d{EP}^{-}}{dt}=p_{{EP}^{+}\to{EP}^{-}}{EP}^{+}-p_{{EP}^{-}\to M}{EP}^{-}$$

(10)

$$\frac{dMLP}{dt}=p_{MPP\to MLP}MPP-p_{MLP\to M}MLP$$

(11)

where $M$ refers to more mature (unmeasured) progenitors. **Eqs. (6)-(11)** apply to both the *Combined* and *Transiting HSCs* models.

We also considered alternative models where MPPs differentiate directly into erythroid progenitors, bypassing their sequential differentiation into CMPs and MEPs. For this, we added a differentiation rate from MPPs to CD71+ EPs, $p_{MPP\to{EP}^{+}}$ and updated **Eqs. (2) and (5)** of both the *Combined* and *Transiting HSCs* models to be:

$$\frac{dMPP}{dt}=p_{HSC\to MPP}HSC-\left( d_{MPP}+p_{MPP\to CMP}+p_{MPP\to MLP}+p_{MPP\to{EP}^{+}} \right)MPP$$

(12)

$$\frac{dMPP}{dt}=p_{HSC1\to MPP}HSC\text{1}+p_{HSC2\to MPP}HSC\text{2}-\left( d_{MPP}+p_{MPP\to CMP}+p_{MPP\to MLP}+p_{MPP\to{EP}^{+}} \right)MPP$$

(13)

with equation (9) rewritten as

$$\frac{d{EP}^{+}}{dt}=p_{MPP\to{EP}^{+}}MPP+p_{MEP\to{EP}^{+}}MEP-\left( d_{{EP}^{+}}+p_{{EP}^{+}\to{EP}^{-}} \right){EP}^{+}.$$

(14)

Finally, for all the hematopoietic hierarchies considered, we also considered MPP self-renewal by simply replacing the outflux rate $-d_{MPP}$ by a positive influx rate caused by net proliferation, $k_{MPP}$. In this case, **Eqs. (2), (5), (12), and (13)** describing MPP dynamics are replaced by

$$\frac{dMPP}{dt}=p_{HSC\to MPP}HSC-\left( p_{MPP\to CMP}+p_{MPP\to MLP}-k_{MPP} \right)MPP,$$

(15)

$$\frac{dMPP}{dt}=p_{HSC\to MPP}HSC-\left( p_{MPP\to CMP}+p_{MPP\to MLP}+p_{MPP\to{EP}^{+}}-k_{MPP} \right)MPP,$$

(16)

$$\frac{dMPP}{dt}=p_{HSC1\to MPP}HSC\text{1}+p_{HSC2\to MPP}HSC\text{2}-\left( p_{MPP\to CMP}+p_{MPP\to MLP}-k_{MPP} \right)MPP,$$

(17)

and

$$\frac{dMPP}{dt}=p_{HSC1\to MPP}HSC\text{1}+p_{HSC2\to MPP}HSC\text{2}-\left( p_{MPP\to CMP}+p_{MPP\to MLP}+p_{MPP\to{EP}^{+}}-k_{MPP} \right)MPP,$$

(18)

respectively.

We solved all eight ODE models in the *DifferentialEquations.jl* package in Julia [1, 2] using the explicit Tsitouras 5/4 Runge-Kutta method via the *Tsit5* algorithm [3].

**Structural identifiability**

A mathematical model is structurally identifiable if it is possible to determine a unique value for each unknown parameter, given measurements of the model’s outputs and its structure [4]. As estimating structurally unidentifiable parameters can lead to incorrect values and misleading results, we performed structural identifiability analysis prior to model calibration using the MATLAB toolbox STRIKE-GOLDD [5]. STRIKE-GOLDD evaluates the change in the output vector (i.e., the observations) along the model’s dynamics by calculating Lie derivatives, which depend on the model’s parameters. If the change in the Lie derivatives leads to linearly independent vectors, we can conclude that at least one parameter is structurally non-identifiable. See **Table S1** for the results of our structural identifiability analysis.

**Parameter estimation**

We calculated Pearson’s correlation coefficient for every pair of parameters within the chosen best fit model for the H3-WT and H3-K27M datasets in Julia using *cor* function from the *Statistics.jl* package [1]. Given that the rates of HSC self-renewal rate and of differentiation into MPPs are highly correlated, we instead estimated the net HSC proliferation rate. Thus, we can rewrite **Eq. (1)** to be

$$\frac{dHSC}{dt}=a_{HSC}HSC,$$

(20)

where $a_{HSC}= k_{HSC}-p_{HSC\to MPP}$. Similarly, Eqs. (3) and (4) become

$$\frac{dHSC\text{1}}{dt}=\left( a_{HSC1}-r_{HSC1\to HSC2} \right)HSC\text{1}+r_{HSC2\to HSC1}HSC\text{2,}$$

(21)

$$\frac{dHSC\text{2}}{dt}=\left( a_{HSC2}-r_{HSC2\to HSC1} \right)HSC\text{2}+r_{HSC1\to HSC2}HSC\text{1},$$

(22)

where $a_{HSC1}= k_{HSC1}-p_{HSC1\to MPP}$ and $a_{HSC2}= k_{HSC2}-p_{HSC2\to MPP}$.

For each model and each condition (H3-WT and H3-K27M), we estimated a parameter set $\boldsymbol{\theta}=(\theta_{1},\ldots,\theta_{n_{p}})$ by fitting the model to cell counts. The number of estimated parameters, $n_{p}$, depends on the hematopoietic hierarchy represented in the mathematical model (i.e., models B, C, D, and E with either the *Combined* or *Transiting* *HSCs* structure; see the Methods in the Main Text). To find best fit values, $\boldsymbol{\theta}^{\boldsymbol{best}}$, we minimized the cost function (i.e., weighted residual sum of squares):

$$WRSS\left( \boldsymbol{\theta} \right)=\sum_{k=1}^{P} \sum_{j=1}^{N} \sum_{i=1}^{M} \frac{({X_{k}^{s}(\boldsymbol{\theta,}t_{i}) -X_{k,j}^{d}(t_{i}))}^{2}}{{\bar{X_{k}}}^{2}}.$$

(19)

Here, $P$ is the number of state variables/blood cell compartments, $X_{k}$ is the $k$th cell population, and $\bar{X_{k}}$ is the mean value of $X_{k}$ over all data points. Let the superscript $s$ refer to the model’s solution and the superscript $d$ refer to a data point. Hence, $X_{k}^{s}(\boldsymbol{\theta,}t_{i})$ is the model’s solution for population $X_{k}$ at time $t_{i}$, given the parameter set $\boldsymbol{\theta}$ and $X_{k}^{d}(t_{i})$ is the cell count of population $X_{k}$ at time $t_{i}$ as measured in our flow cytometry data. Lastly, $M=4$ is the number of measured time points, and $N=9$ is the number of mice replicates at each time point.

The best fit value, $\boldsymbol{\theta}^{\boldsymbol{best}}$, corresponds to the global minimum of the cost function (**Eq. (19)**), which we found using Ingber’s adaptive simulated annealing (ASA) [6, 7]. When considering parameters with varying finite ranges (i.e., bounded by real physical or biological properties), the annealing schedule in ASA is much faster than the one used in the standard simulated annealing algorithm. ASA also take into consideration that the cost function has different sensitivities to each parameter, allowing for optimizing annealing time for a faster convergence of the algorithm. Briefly, our ASA algorithm works as follows.

1. An initial set of parameters, $\boldsymbol{\theta}^{\boldsymbol{old}}$, is randomly selected, with each parameter guess, $\theta_{i}^{old}$, $i\in[1,...,n_{p}]$, chosen uniformly between their given lower- and upper-bound [$L_{i},U_{i}$] values. Then, we initialize the algorithm by setting the initial generating annealing times, $n_{i}=0$ , $i\in[1,...,n_{p}]$, and the acceptance annealing time, $n_{a}=0$. We also set the initial generating temperatures $T_{i}\left( 0 \right)=1.0$, $i\in[1,...,n_{p}]$, and the acceptance temperature $T_{a}\left( 0 \right)=WRSS\left( \boldsymbol{\theta}^{\boldsymbol{old}} \right)$. We start the iteration with one generated and one accepted parameter set, so that $N_{gen}=1$ and $N_{acc}=1$, respectively.
2. We randomly generate a new point, $\boldsymbol{\theta}^{\boldsymbol{new}}$, per the following equation:

$\theta_{i}^{new}=\theta_{i}^{old}+g_{i}\cdot(U_{i}-L_{i})$, $i\in[1,...,n_{p}],$

where

$$g_{i}=sgn(u_{i}-\frac{1}{2})\cdot T_{i}(n_{i})\cdot\left[ \left( 1+\frac{1}{T_{i}\left( n_{i} \right)} \right)^{\left| 2u_{i}-1 \right|}-1 \right].$$

In the expression of $g_{i}$ above, $u_{i}\sim\mathcal{U}\left( 0,1 \right)$, $i\in[1,...,n_{p}]$, is a uniformly distributed random variable. We discard $\boldsymbol{\theta}^{\boldsymbol{new}}$ if any $\theta_{i}^{new}$ lies outside the parameter’s bounds [$L_{i},U_{i}$]. Step 2 is repeated until otherwise and a new point is generated. We update $N_{gen}=N_{gen}+1$.

3. If $WRSS(\boldsymbol{\theta}^{\boldsymbol{new}})<WRSS(\boldsymbol{\theta}^{\boldsymbol{old}})$, then $\boldsymbol{\theta}^{\boldsymbol{new}}$ is accepted. Otherwise, we calculate the acceptance probability of $\boldsymbol{\theta}^{\boldsymbol{new}}$ given by the standard Metropolis acceptance criterion:

$$P_{acc}=exp\left( -\frac{WRSS\left( \boldsymbol{\theta}^{\boldsymbol{new}} \right)-WRSS(\boldsymbol{\theta}^{\boldsymbol{old}})}{T_{a}(n_{a})} \right)$$

and generate a random variable $q\sim\mathcal{U}\left( 0,1 \right)$. If$q<P_{acc}$, then $\boldsymbol{\theta}^{\boldsymbol{new}}$ is accepted. If not, it is rejected. When $\boldsymbol{\theta}^{\boldsymbol{new}}$ is accepted we update $N_{acc}=N_{acc}+1$ and $\boldsymbol{\theta}^{\boldsymbol{old}}=\boldsymbol{\theta}^{\boldsymbol{new}}$.

4. Temperature annealing occurs when $N_{gen}$ gets sufficiently large. We set $N_{gen}=800$. For all $i\in[1,...,n_{p}]$, we update $n_{i}=n_{i}+1$ and calculate

${T_{i}\left( n_{i} \right)=T}_{i}\left( 0 \right)\cdot exp\left( -m{n_{i}}^{\frac{1}{n_{p}}} \right), i\in[1,...,n_{p}]$.

Similarly, we change $n_{a}=n_{a}+1$ and set

${T_{a}\left( n_{a} \right)=T}_{a}\left( 0 \right)\cdot exp\left( -m{n_{a}}^{\frac{1}{n_{p}}} \right)$.

where $m \text{∈}\text{ [}1,10\text{]}$ is chosen; in our case, we took $m=$ $5$. At the end of the temperature annealing step, we reinitialize $N_{gen}=1$.

5. Reannealing takes place when $N_{acc}$ is sufficiently large. We chose $N_{acc}=30$ to optimize both computation time and acceptance ratio (i.e., the total number of accepted points over the total number of cost function evaluations). During this step, we scale the generating temperatures, $T_{i}$, with the sensitivity of $WRSS(\boldsymbol{\theta})$ to each $\theta_{i}$, $i\in[1,...,n_{p}]$, given by

$$s_{i}=\left| \frac{WRSS\left( \boldsymbol{\theta}^{\boldsymbol{best}}+\varepsilon\theta_{i}^{best}\boldsymbol{e}_{\boldsymbol{i}} \right))-WRSS(\boldsymbol{\theta}^{\boldsymbol{best}})}{\varepsilon\theta_{i}^{best}} \right|,$$

where $\varepsilon=0.005$ and $\boldsymbol{e}_{\boldsymbol{i}}$ is the $n_{p}$-dimensional base vector. We chose to calculate a relative perturbation to measure a sensitivity that is consistent across parameters with varying magnitudes. Thus, we reset $T_{i}\left( 0 \right)=1.0$ and update $T_{i}\left( n_{i} \right)=\frac{s_{max}}{s_{i}}T_{i}\left( n_{i} \right)$ for all $i\in[1,...,n_{p}]$, where $s_{max}$ is the highest element of all $s_{i}$, $i\in[1,...,n_{p}]$. Similarly, we reset $T_{a}(0)$ to the last accepted $WRSS(\boldsymbol{\theta}^{\boldsymbol{new}})$ and $T_{a}\left( n_{a} \right)=WRSS(\boldsymbol{\theta}^{\boldsymbol{best}})$. Furthermore, we rescale the annealing times accordingly:

$$n_{i}=\left( -\frac{1}{m}\cdot log\left( \frac{T_{i}\left( n_{i} \right)}{T_{i}\left( 0 \right)} \right) \right)^{n_{p}}, i\in[1,...,n_{p}]$$

$$n_{a}=\left( -\frac{1}{m}\cdot log\left( \frac{T_{a}\left( n_{a} \right)}{T_{a}\left( 0 \right)} \right) \right)^{n_{p}}.$$

At the end of the step, we reinitialize $N_{acc}=1$.

6. We repeat steps 2-5 until $WRSS\left( \boldsymbol{\theta}^{\boldsymbol{best}} \right)$remains within a small tolerance, $\delta_{C}$, for $k$ reannealing steps or meets the maximum amount of cost function evaluations, $e_{max}$. In this work, we chose $\delta_{C}=0.05$, $k=5$, and $e_{max}=5\times{10}^{5}$. Once these tolerances are reached, the algorithm ends.

We implemented our ASA algorithm in Julia, using the *Distributions.jl* package [8] to generate the random variables with the standard uniform distribution for the creation of new points in Step 2. Due to the random nature of ASA, we performed 200 iterations of the global optimization algorithm to get the best fit parameter set for all models. We then selected from the 200 iteration samples the parameter set $\boldsymbol{\theta}$ that corresponds to the minimum the cost function value over these 200 iterations.

**Distributions and credible intervals of best fit estimates**

Distributions for the best fit parameter values of the H3-WT and H3-K27M selected models were defined using trajectory-matching random-walk feasibility sampling (TM-RWFS) [9] (see **Methods**; **Figure 3A;** **Figure S5**). The resulting trajectories for the selected models are given in **Figure S4**. We measured the 95% credible interval from those distributions by calculating the 95% highest density interval (HDI) [10]. These calculations were performed in Julia by using the function *hdi* from the *PosteriorStats.jl* package [11].

**Statistical analysis of candidate mathematical models**

To find out which model best represents our data, we calculated the corrected Akaike information criteria (AICc) for each candidate model after fitting to both H3-WT and H3-K27M cell counts. The AICc is given by

$$AICc=N\cdot\text{ln}\left( \frac{WRSS(\boldsymbol{\theta}^{\boldsymbol{best}}\mathbf{)}}{N} \right)+\frac{2n_{p}N}{N-n_{p}-1},$$

(23)

with $n_{p}$ and $N$ defined as above. Since $n_{p}\geq N/40$. Here, $WRSS\left( \boldsymbol{\theta}^{\boldsymbol{best}} \right)$, calculated from **Eq. (19)**, is the lowest cost function result from the ASA algorithm samples. For better interpretation, we calculated the $\text{Δ}\mathrm{AICc}$ score given by the difference between the best model score to the other models’ score, i.e.,

$$\text{Δ}{AICc}_{i}={AICc}_{i}-\min_{i\in[1, \ldots, M]} AICc$$

(24)

for $M$ different models. The models with the best expected predictive performance for H3-WT and H3-K27M have $\text{Δ}AICc=0$, i.e., the minimum AICc value, given their best fit values to the data.

**Statistical analysis**

Unequal variance two-sample t-test

To test whether the cell counts at day 112 of each hematopoietic population are statistically different between the H3-WT and H3-K27M genotypes, we performed an unequal variance two-sample t-test (or Welch’s t-test) [12]. We evaluated the null hypothesis that both are independent random samples from normal distributions with equal means, without assuming equal variance. We implemented it in Julia [1], using the function *UnequalVarianceTTest* from *HypothesisTests.jl* package [13].

Mann-Whitney U test

The Mann-Whitney U test measures whether two random variables, such as the best fit values for the H3-WT and H3-K27M models, have the same distributions. This is a nonparametric alternative to the *t*-test when a normal distribution cannot be assumed for the distribution of the random variable. The rank-biserial correlation ($r_{rb}$) is an effect size measure for the Mann–Whitney U test that ranges from −1 to +1. An $r_{rb}$ of 0 indicates no association between group membership and the outcome, $r_{rb}>0$ indicates that the first group tends to have higher ranks, and $r_{rb}<0$ indicates that the second group tends to have higher ranks. In general, larger absolute values denote stronger effects.

To perform the Mann-Whitney U test, we first consolidate the values from the two random variable distributions into a single list. Then we assign sequential ranks in ascending order. If two values are equal, they are given the same rank. Let $n_{1}$ and $n_{2}$ be the number of observations and $R_{1}$and $R_{2}$ be the sum of the ranks for the first and second distributions, respectively. The U statistic for $i=1,2$ is

$$U_{i}=R_{i}-\frac{n_{i}(n_{i}+1)}{2}.$$

We then take $U= \min(U_{1},U_{2})$. Then, the rank-biserial correlation is given by

$$r_{rb}=1-\frac{2U}{n_{1}n_{2}}.$$

**Normalization of cell count solutions**

To better compare the solutions of the H3-K27M fit model to those from wild-type, we calculated the normalized cell count. This was done in two ways: by dividing each H3-WT and H3-K27M cell populations by their initial value at t = 28 days (**Figure S4A**) and by taking the ratio of H3-K27M over H3-WT for all cell compartments (**Figure S4B**). For the two conditions, we compare the entire HSC population by taking the sum of the solutions for H3-K27M HSC1 and HSC2 subpopulations.

### Supplementary Information

**Table S1. Structural identifiability and observability analysis for models B, C, D, and E with the Combined HSCs and Transiting HSCs.** Related to Figure 1 in the Main Text.

| Model | No. states | No.  parameters | Rank | Observable states | Indentifiable parameters | Overall result |
| --- | --- | --- | --- | --- | --- | --- |
| B; Combined HSCs | 8 | 15 | 23 | 8/8 | 15/15 | All states observable; all parameters identifiable |
| C; Combined HSCs | 8 | 15 | 23 | 8/8 | 15/15 | All states observable; all parameters identifiable |
| D; Combined HSCs | 8 | 16 | 24 | 8/8 | 16/16 | All states observable; all parameters identifiable |
| E; Combined HSCs | 8 | 16 | 24 | 8/8 | 16/16 | All states observable; all parameters identifiable |
| B; Transiting HSCs | 9 | 19 | 28 | 9/9 | 19/19 | All states observable; all parameters identifiable |
| C; Transiting HSCs | 9 | 19 | 28 | 9/9 | 19/19 | All states observable; all parameters identifiable |
| D; Transiting HSCs | 9 | 20 | 29 | 9/9 | 20/20 | All states observable; all parameters identifiable |
| E; Transiting HSCs | 9 | 20 | 29 | 9/9 | 20/20 | All states observable; all parameters identifiable |

**Table S2. Estimated parameter values and 95% credible intervals for selected model C with the Combined HSCs (H3-WT) and Transiting HSCs (H3-K27M) structures.** Related to Figure 1 in the Main Text.

|  | **H3-WT** | | | **H3-K27M** | | |
| --- | --- | --- | --- | --- | --- | --- |
|  | *Model C with Combined HSCs* | | | *Model C with Transiting HSCs* | | |
| **Biological Role** | **Parameter** | **Best fit value (days^-1^)** | **95% CI** | **Parameter** | **Best fit value (days^-1^)** | **95% CI** |
| Proliferation | $a_{HSC}$* | 0.017 | [1.3x10^-3^, 0.020] | $a_{HSC1}$** | 0.031 | [1.5x10^-4^, 0.038] |
|  |  |  |  | $a_{HSC2}$*** | 0.011 | [6.3x10^-5^, 0.042] |
|  | $k_{MPP}$ | 5.016 | [4.787, 5.754] | $k_{MPP}$ | 3.856 | [3.445, 4.396] |
| Differentiation | $p_{HSC\to MPP}$ | 0.400 | [0.132, 0.500] | $p_{HSC1\to MPP}$ | 0.467 | [0.045, 0.469] |
|  |  |  |  | $p_{HSC2\to MPP}$ | 0.135 | [0.057, 0.497] |
|  | $p_{MPP\to MLP}$ | 0.092 | [0.024, 0.100] | $p_{MPP\to MLP}$ | 0.063 | [0.012, 0.069] |
|  | $p_{MPP\to CMP}$ | 5.365 | [4.962, 5.931] | $p_{MPP\to CMP}$ | 4.715 | [4.064, 5.266] |
|  | $p_{CMP\to GMP}$ | 0.097 | [0.064, 0.199] | $p_{CMP\to GMP}$ | 0.030 | [0.011, 0.060] |
|  | $p_{CMP\to MEP}$ | 1.451 | [1.032, 1.852] | $p_{CMP\to MEP}$ | 2.190 | [1.094, 2.500] |
|  | $p_{MEP\to EP+}$ | 4.619 | [4.189, 5.845] | $p_{MEP\to EP+}$ | 6.382 | [4.782, 6.999] |
|  | $p_{EP+\to EP-}$ | 1.122 | [0.571, 2.899] | $p_{EP+\to EP-}$ | 0.411 | [0.016, 1.156] |
|  | $p_{MLP\to M}$ | 0.026 | [2.5x10^-5^, 0.079] | $p_{MLP\to M}$ | 0.106 | [1.7x10^-3^, 0.183] |
|  | $p_{GMP\to M}$ | 3.1x10^-3^ | [2.2x10^-6^, 0.047] | $p_{GMP\to M}$ | 1.0x10^-3^ | [2.7x10^-4^, 9.6x10^-3^] |
|  | $p_{EP-\to M}$ | 0.748 | [0.434,0.958] | $p_{EP-\to M}$ | 1.652 | [0.037, 2.119] |
| Death | $d_{CMP}$ | 0.288 | [0.014, 0.982] | $d_{CMP}$ | 0.344 | [0.120, 0.989] |
|  | $d_{MEP}$ | 0.092 | [3.9x10^-4^, 0.656] | $d_{MEP}$ | 0.033 | [0.018, 0.199] |
|  | $d_{EP+}$ | 0.249 | [0.066,1.919] | $d_{EP+}$ | 0.358 | [0.031, 0.830] |
| Transition | N/A | | | $r_{HSC1\to HSC2}$ | 0.021 | [2.0x10^-4^, 0.040] |
|  |  |  |  | $r_{HSC2\to HSC1}$ | 0.017 | [7.8x10^-4^, 0.085] |

***** $a_{HSC}=k_{HSC}-p_{HSC\to MPP}$

****** $a_{HSC1}=k_{HSC1}-p_{HSC1\to MPP}$

******* $a_{HSC}=k_{HSC2}-p_{HSC2\to MPP}$

**Table S3. Estimated parameter values for models B, D, and E with Combined HSCs structure for H3-WT.** Related to Figure 1 in the Main Text.

| **H3-WT** | | | | | |
| --- | --- | --- | --- | --- | --- |
| *Combined HSCs* | | | | | |
| **Model B** | | **Model D** | | **Model E** | |
| **Parameter** | **Best fit value (days^-1^)** | **Parameter** | **Best fit value (days^-1^)** | **Parameter** | **Best fit value (days^-1^)** |
| $a_{HSC}$* | 0.016 | $a_{HSC}*$ | 0.019 | $a_{HSC}*$ | 0.018 |
|  |  |  |  | $k_{MPP}$ | 5.327 |
| $p_{HSC\to MPP}$ | 5.458 | $p_{HSC\to MPP}$ | 6.000 | $p_{HSC\to MPP}$ | 0.364 |
| $p_{MPP\to MLP}$ | 0.168 | $p_{MPP\to MLP}$ | 0.130 | $p_{MPP\to MLP}$ | 0.089 |
| $p_{MPP\to CMP}$ | 3.936 | $p_{MPP\to CMP}$ | 5.348 | $p_{MPP\to CMP}$ | 4.345 |
|  |  | $p_{MPP\to EP+}$ | 3.736 | $p_{MPP\to EP+}$ | 1.209 |
| $p_{CMP\to GMP}$ | 0.193 | $p_{CMP\to GMP}$ | 0.829 | $p_{CMP\to GMP}$ | 0.240 |
| $p_{CMP\to MEP}$ | 0.972 | $p_{CMP\to MEP}$ | 1.342 | $p_{CMP\to MEP}$ | 1.589 |
| $p_{MEP\to EP+}$ | 1.524 | $p_{MEP\to EP+}$ | 0.884 | $p_{MEP\to EP+}$ | 1.962 |
| $p_{EP+\to EP-}$ | 0.293 | $p_{EP+\to EP-}$ | 1.117 | $p_{EP+\to EP-}$ | 0.852 |
| $p_{MLP\to M}$ | 0.062 | $p_{MLP\to M}$ | 0.041 | $p_{MLP\to M}$ | 0.048 |
| $p_{GMP\to M}$ | 0.021 | $p_{GMP\to M}$ | 0.058 | $p_{GMP\to M}$ | 0.026 |
| $p_{EP-\to M}$ | 0.209 | $p_{EP-\to M}$ | 0.563 | $p_{EP-\to M}$ | 0.589 |
| $d_{MPP}$ | 1.600 | $d_{MPP}$ | 0.697 |  |  |
| $d_{CMP}$ | 0.268 | $d_{CMP}$ | 0.957 | $d_{CMP}$ | 0.193 |
| $d_{MEP}$ | 1.502 | $d_{MEP}$ | 1.432 | $d_{MEP}$ | 2.731 |
| $d_{EP+}$ | 0.130 | $d_{EP+}$ | 0.339 | $d_{EP+}$ | 0.387 |

***** $a_{HSC}=k_{HSC}-p_{HSC\to MPP}$

**Table S4. Estimated parameter values for models B, C, D, and E with Transiting HSCs structure for H3-WT.** Related to Figure 1 in the Main Text.

| **H3-WT** | | | | | | | |
| --- | --- | --- | --- | --- | --- | --- | --- |
| *Transiting HSCs* | | | | | | | |
| **Model B** | | **Model C** | | **Model D** | | **Model E** | |
| **Parameter** | **Best fit value (days^-1^)** | **Parameter** | **Best fit value (days^-1^)** | **Parameter** | **Best fit value (days^-1^)** | **Parameter** | **Best fit value (days^-1^)** |
| $a_{HSC1}$* | 1.6x10^-3^ | $a_{HSC1}$* | 9.6x10^-4^ | $a_{HSC1}$* | 1.0x10^-4^ | $a_{HSC1}$* | 2.1x10^-3^ |
| $a_{HSC2}$** | 0.020 | $a_{HSC2}$** | 0.021 | $a_{HSC2}$** | 0.016 | $a_{HSC2}$** | 0.019 |
|  |  | $k_{MPP}$ | 5.971 |  |  | $k_{MPP}$ | 5.823 |
| $p_{HSC1\to MPP}$ | 7.255 | $p_{HSC1\to MPP}$ | 0.457 | $p_{HSC1\to MPP}$ | 7.230 | $p_{HSC1\to MPP}$ | 0.465 |
| $p_{HSC2\to MPP}$ | 7.860 | $p_{HSC2\to MPP}$ | 0.398 | $p_{HSC2\to MPP}$ | 5.436 | $p_{HSC2\to MPP}$ | 0.228 |
| $p_{MPP\to MLP}$ | 0.169 | $p_{MPP\to MLP}$ | 0.096 | $p_{MPP\to MLP}$ | 0.148 | $p_{MPP\to MLP}$ | 0.110 |
| $p_{MPP\to CMP}$ | 7.495 | $p_{MPP\to CMP}$ | 6.280 | $p_{MPP\to CMP}$ | 5.050 | $p_{MPP\to CMP}$ | 4.567 |
|  |  |  |  | $p_{MPP\to EP+}$ | 0.870 | $p_{MPP\to EP+}$ | 1.453 |
| $p_{CMP\to GMP}$ | 1.351 | $p_{CMP\to GMP}$ | 0.789 | $p_{CMP\to GMP}$ | 0.381 | $p_{CMP\to GMP}$ | 0.339 |
| $p_{CMP\to MEP}$ | 1.175 | $p_{CMP\to MEP}$ | 1.508 | $p_{CMP\to MEP}$ | 1.528 | $p_{CMP\to MEP}$ | 1.253 |
| $p_{MEP\to EP+}$ | 3.496 | $p_{MEP\to EP+}$ | 4.397 | $p_{MEP\to EP+}$ | 1.955 | $p_{MEP\to EP+}$ | 0.954 |
| $p_{EP+\to EP-}$ | 0.852 | $p_{EP+\to EP-}$ | 0.877 | $p_{EP+\to EP-}$ | 0.648 | $p_{EP+\to EP-}$ | 0.574 |
| $p_{MLP\to M}$ | 0.056 | $p_{MLP\to M}$ | 0.037 | $p_{MLP\to M}$ | 0.062 | $p_{MLP\to M}$ | 0.036 |
| $p_{GMP\to M}$ | 0.211 | $p_{GMP\to M}$ | 0.124 | $p_{GMP\to M}$ | 0.056 | $p_{GMP\to M}$ | 0.047 |
| $p_{EP-\to M}$ | 0.640 | $p_{EP-\to M}$ | 0.636 | $p_{EP-\to M}$ | 0.453 | $p_{EP-\to M}$ | 0.386 |
| $d_{MPP}$ | 1.396 |  |  | $d_{MPP}$ | 0.009 |  |  |
| $d_{CMP}$ | 0.195 | $d_{CMP}$ | 0.107 | $d_{CMP}$ | 0.040 | $d_{CMP}$ | 0.098 |
| $d_{MEP}$ | 0.117 | $d_{MEP}$ | 0.647 | $d_{MEP}$ | 2.587 | $d_{MEP}$ | 2.917 |
| $d_{EP+}$ | 0.082 | $d_{EP+}$ | 0.213 | $d_{EP+}$ | 0.192 | $d_{EP+}$ | 0.174 |
| $r_{HSC1\to HSC2}$ | 3.7x10^-3^ | $r_{HSC1\to HSC2}$ | 1.0x10^-3^ | $r_{HSC1\to HSC2}$ | 9.1x10^-3^ | $r_{HSC1\to HSC2}$ | 4.6x10^-3^ |
| $r_{HSC2\to HSC1}$ | 4.0x10^-3^ | $r_{HSC2\to HSC1}$ | 3.3x10^-3^ | $r_{HSC2\to HSC1}$ | 4.7x10^-3^ | $r_{HSC2\to HSC1}$ | 3.4x10^-3^ |

***** $a_{HSC1}=k_{HSC1}-p_{HSC1\to MPP}$

****** $a_{HSC}=k_{HSC2}-p_{HSC2\to MPP}$

**Table S5. Estimated parameter values for models B, C, D, and E with the Combined HSCs structure for H3-K27M.** Related to Figure 1 in the Main Text.

| **H3-K27M** | | | | | | | |
| --- | --- | --- | --- | --- | --- | --- | --- |
| *Combined HSCs* | | | | | | | |
| **Model B** | | **Model C** | | **Model D** | | **Model E** | |
| **Parameter** | **Best fit value (days^-1^)** | **Parameter** | **Best fit value (days^-1^)** | **Parameter** | **Best fit value (days^-1^)** | **Parameter** | **Best fit value (days^-1^)** |
| $a_{HSC}$* | 0.044 | $a_{HSC}$* | 0.044 | $a_{HSC}$* | 0.044 | $a_{HSC}$* | 0.044 |
|  |  | $k_{MPP}$ | 4.438 |  |  | $k_{MPP}$ | 2.516 |
| $p_{HSC\to MPP}$ | 3.961 | $p_{HSC\to MPP}$ | 0.496 | $p_{HSC\to MPP}$ | 3.712 | $p_{HSC\to MPP}$ | 0.468 |
| $p_{MPP\to MLP}$ | 0.021 | $p_{MPP\to MLP}$ | 0.042 | $p_{MPP\to MLP}$ | 0.095 | $p_{MPP\to MLP}$ | 0.034 |
| $p_{MPP\to CMP}$ | 6.278 | $p_{MPP\to CMP}$ | 5.763 | $p_{MPP\to CMP}$ | 7.081 | $p_{MPP\to CMP}$ | 3.231 |
|  |  |  |  | $p_{MPP\to EP+}$ | 2.109 | $p_{MPP\to EP+}$ | 0.613 |
| $p_{CMP\to GMP}$ | 0.839 | $p_{CMP\to GMP}$ | 0.025 | $p_{CMP\to GMP}$ | 1.529 | $p_{CMP\to GMP}$ | 0.026 |
| $p_{CMP\to MEP}$ | 1.894 | $p_{CMP\to MEP}$ | 2.696 | $p_{CMP\to MEP}$ | 1.800 | $p_{CMP\to MEP}$ | 1.238 |
| $p_{MEP\to EP+}$ | 5.637 | $p_{MEP\to EP+}$ | 7.876 | $p_{MEP\to EP+}$ | 1.790 | $p_{MEP\to EP+}$ | 3.011 |
| $p_{EP+\to EP-}$ | 0.470 | $p_{EP+\to EP-}$ | 0.471 | $p_{EP+\to EP-}$ | 0.352 | $p_{EP+\to EP-}$ | 0.388 |
| $p_{MLP\to M}$ | 3.6x10^-3^ | $p_{MLP\to M}$ | 0.048 | $p_{MLP\to M}$ | 0.159 | $p_{MLP\to M}$ | 0.027 |
| $p_{GMP\to M}$ | 1.109 | $p_{GMP\to M}$ | 5.6x10^-4^ | $p_{GMP\to M}$ | 2.049 | $p_{GMP\to M}$ | 1.6x10^-4^ |
| $p_{EP-\to M}$ | 1.616 | $p_{EP-\to M}$ | 1.734 | $p_{EP-\to M}$ | 1.242 | $p_{EP-\to M}$ | 1.204 |
| $d_{MPP}$ | 4.832 |  |  | $d_{MPP}$ | 1.700 |  |  |
| $d_{CMP}$ | 0.464 | $d_{CMP}$ | 0.260 | $d_{CMP}$ | 0.258 | $d_{CMP}$ | 0.194 |
| $d_{MEP}$ | 0.331 | $d_{MEP}$ | 0.331 | $d_{MEP}$ | 4.121 | $d_{MEP}$ | 0.696 |
| $d_{EP+}$ | 0.238 | $d_{EP+}$ | 0.529 | $d_{EP+}$ | 0.272 | $d_{EP+}$ | 0.148 |

***** $a_{HSC}=k_{HSC}-p_{HSC\to MPP}$

**Table S6. Estimated parameter values for models B, D, and E with the Transiting HSCs structure for H3-K27M.** Related to Figure 1 in the Main Text.

| **H3-K27M** | | | | | |
| --- | --- | --- | --- | --- | --- |
| *Transiting HSCs* | | | | | |
| **Model B** | | **Model D** | | **Model E** | |
| **Parameter** | **Best fit value (days^-1^)** | **Parameter** | **Best fit value (days^-1^)** | **Parameter** | **Best fit value (days^-1^)** |
| $a_{HSC1}$* | 0.039 | $a_{HSC1}$* | 0.040 | $a_{HSC1}$* | 0.035 |
| $a_{HSC2}$** | 0.044 | $a_{HSC2}$** | 0.021 | $a_{HSC2}$** | 0.031 |
|  |  |  |  | $k_{MPP}$ | 5.681 |
| $p_{HSC1\to MPP}$ | 4.568 | $p_{HSC1\to MPP}$ | 6.392 | $p_{HSC1\to MPP}$ | 0.442 |
| $p_{HSC2\to MPP}$ | 1.176 | $p_{HSC2\to MPP}$ | 0.178 | $p_{HSC2\to MPP}$ | 0.312 |
| $p_{MPP\to MLP}$ | 0.030 | $p_{MPP\to MLP}$ | 0.056 | $p_{MPP\to MLP}$ | 0.061 |
| $p_{MPP\to CMP}$ | 7.566 | $p_{MPP\to CMP}$ | 7.908 | $p_{MPP\to CMP}$ | 4.484 |
|  |  | $p_{MPP\to EP+}$ | 3.628 | $p_{MPP\to EP+}$ | 2.214 |
| $p_{CMP\to GMP}$ | 1.362 | $p_{CMP\to GMP}$ | 0.377 | $p_{CMP\to GMP}$ | 0.033 |
| $p_{CMP\to MEP}$ | 2.252 | $p_{CMP\to MEP}$ | 3.200 | $p_{CMP\to MEP}$ | 2.051 |
| $p_{MEP\to EP+}$ | 5.952 | $p_{MEP\to EP+}$ | 8.491 | $p_{MEP\to EP+}$ | 6.787 |
| $p_{EP+\to EP-}$ | 0.536 | $p_{EP+\to EP-}$ | 1.016 | $p_{EP+\to EP-}$ | 0.631 |
| $p_{MLP\to M}$ | 0.024 | $p_{MLP\to M}$ | 0.077 | $p_{MLP\to M}$ | 0.105 |
| $p_{GMP\to M}$ | 1.797 | $p_{GMP\to M}$ | 0.444 | $p_{GMP\to M}$ | 4.2x10^-3^ |
| $p_{EP-\to M}$ | 1.833 | $p_{EP-\to M}$ | 3.695 | $p_{EP-\to M}$ | 2.547 |
| $d_{MPP}$ | 2.126 | $d_{MPP}$ | 0.576 |  |  |
| $d_{CMP}$ | 0.265 | $d_{CMP}$ | 0.461 | $d_{CMP}$ | 0.289 |
| $d_{MEP}$ | 1.140 | $d_{MEP}$ | 1.436 | $d_{MEP}$ | 0.215 |
| $d_{EP+}$ | 0.220 | $d_{EP+}$ | 0.739 | $d_{EP+}$ | 0.539 |
| $r_{HSC1\to HSC2}$ | 3.0x10^-3^ | $r_{HSC1\to HSC2}$ | 0.014 | $r_{HSC1\to HSC2}$ | 8.7x10^-3^ |
| $r_{HSC2\to HSC1}$ | 6.2x10^-3^ | $r_{HSC2\to HSC1}$ | 1.9x10^-3^ | $r_{HSC2\to HSC1}$ | 0.015 |

***** $a_{HSC1}=k_{HSC1}-p_{HSC1\to MPP}$

****** $a_{HSC}=k_{HSC2}-p_{HSC2\to MPP}$

**Table S7. Results of Mann-Whitney U tests of the distributions of H3-K27M compared to those of H3-WT for the shared parameters in the selected model C with the Combined HSCs and Transiting HSCs structure.** Related to Figure 1 in the Main Text. Results for parameter estimation to H3-WT and H3-K27M experimental data.

| **Biological Role** | **Parameter** | **Effect Size (%)** | **Rank-biserial**  **Correlation (**$\boldsymbol{r}_{\boldsymbol{rb}}$**)** | **p-value** |
| --- | --- | --- | --- | --- |
| Proliferation | $k_{MPP}$ | 0 | -1 | 0 |
| Differentiation | $p_{MPP\to MLP}$ | 17.92 | -0.642 | < 1x10^-16^ |
|  | $p_{MPP\to CMP}$ | 2.88 | -0.942 | < 1x10^-16^ |
|  | $p_{CMP\to GMP}$ | 0.60 | -0.988 | < 1x10^-16^ |
|  | $p_{CMP\to MEP}$ | 74.72 | 0.494 | < 1x10^-16^ |
|  | $p_{MEP\to EP+}$ | 88.43 | 0.769 | < 1x10^-16^ |
|  | $p_{EP+\to EP-}$ | 7.17 | -0.856 | < 1x10^-16^ |
|  | $p_{MLP\to M}$ | 83.41 | 0.668 | < 1x10^-16^ |
|  | $p_{GMP\to M}$ | 14.38 | -0.712 | < 1x10^-16^ |
|  | $p_{EP-\to M}$ | 68.07 | 0.361 | < 1x10^-16^ |
| Death | $d_{CMP}$ | 75.00 | 0.500 | < 1x10^-16^ |
|  | $d_{MEP}$ | 20.55 | -0.589 | < 1x10^-16^ |
|  | $d_{EP+}$ | 26.94 | -0.461 | < 1x10^-16^ |

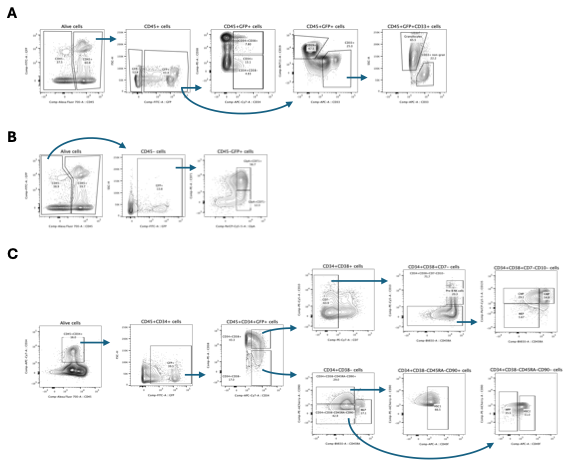

**Figure S1. Gating strategy for the analyzed cell populations across three flow cytometry panels, using representative plots from week-16 H3-K27M samples.** Global panel **(A)** and differentiation panel **(B)** were both performed on unenriched cell populations. HSC panel **(C)** was performed following the depletion of murine cells and human lineage-committed cells.

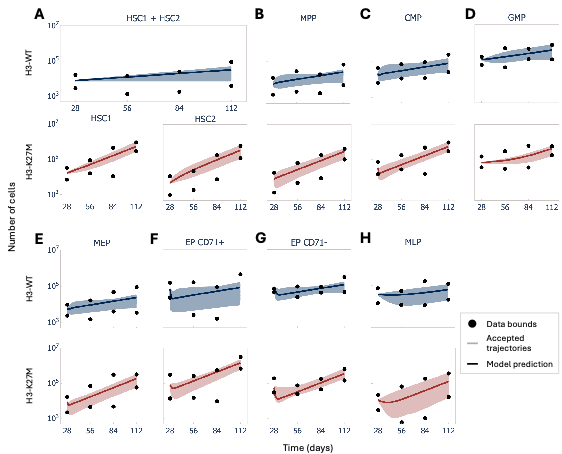

**Figure S2. Output of TM-RWFS method on selected hematopoietic hierarchy models.** Accepted trajectories (N = 1000) for model C for H3-WT (blue; Combined HSCs structure) and H3-K27M (red; Transiting HSCs structure) cells across hematopoietic populations. **A)** HSC1+HSC2 combined for H3-WT and HSC1 and HSC2 for H3-K27M. **B)** MPP. **C)** CMP. **D)** GMP. **E)** MEP. **F)** EP CD71+. **G)** EP CD71−. **H)** MLP. Solid circles indicate data bounds for each time point; pale solid lines indicate accepted trajectories; dark solid lines indicate model prediction.

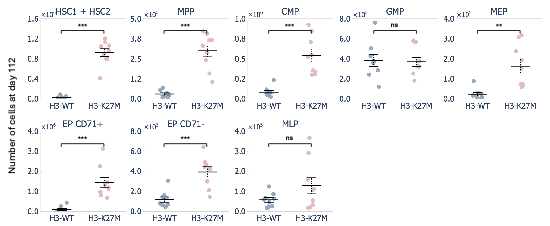

**Figure S3. H3-K27M cell counts are significantly higher at day 112 after transplantation in all hematopoietic populations, except GMPs and MPPs.** Mean cell count values at day 112 between H3-WT and H3-K27M were compared with an unequal variance two-sample t-test (see Supplementary Methods). Solid circles describe cell counts for H3-WT (blue) and H3-K27M (red); tick solid lines represent mean values; error bars correspond to standard errors. Ns: p > 0.05, **: p ≤ 0.01, ***: p ≤ 0.001.

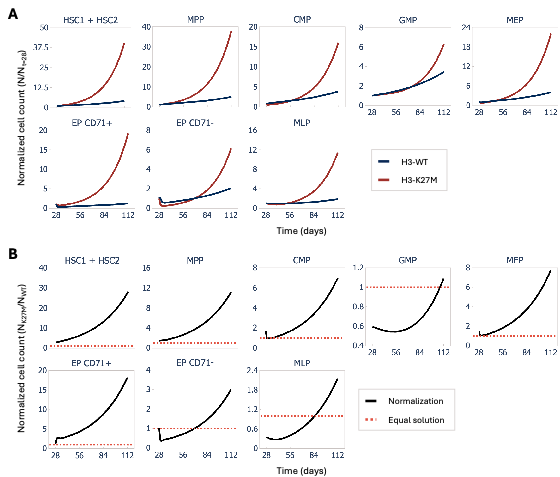

**Figure S4. H3-K27M increases the expansion of all blood cell populations compared to wild-type.** A) Results of predictions from Model C using the Combined (H3-WT) or Transiting HSC (H3-K27M) structure, normalized to the initial number of cells (t = 28 days). Blue solid lines: H3-WT. Red solid lines: H3-K27M. B) Results of predictions from Model C with Transiting HSC (H3-K27M) structure normalized to predictions from Model C with Combined HSC (H3-WT) structure. Red horizontal dotted line**s** at 1 show where solutions are equal. Below the line, there are more cells for H3-WT than H3-K27M. Above the line, H3-K27M has a larger cell count than wild-type. Related to Figure 2 in the Main Text.

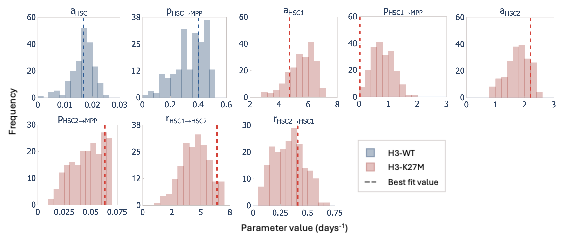

**Figure S5. Distribution of model parameters from the HSC compartments.** Distributions of parameter estimates based on 1,000 accepted trajectories using TM-RWFS. Each subplot displays the distributions of a parameter from the HSC compartment of model C fit to H3-WT (blue; Combined HSCs structure) and H3-K27M (red; Transiting HSCs structure) data. Vertical dashed lines represent best fit values.
